# Towards Understanding the Drivers of Antibody-Antigen Binding

**DOI:** 10.64898/2026.01.04.697575

**Authors:** Omeir Khan, Marcus Kankkunen, Michel Shaker, Henry Chow, Nikolas Moustakas, Siri Allegra-Berger, Sandor Vajda, Diane Joseph-McCarthy

## Abstract

Antibody therapeutics are capable of binding to target antigens with a high degree of specificity and affinity. Gaining an understanding of how antibody–antigen interactions are governed can provide valuable insights that may assist with rational paratope design and epitope prediction. In this work, we apply the FTMap algorithm to systematically characterize binding hot spots – regions on a protein surface that contribute disproportionately to molecular recognition, to a set of 50 antibody–antigen complexes. From our analysis, we find that interface hot spots are typically concentrated on the paratope (antibody side) of the interface, indicating that paratopes typically function as hot spot rich environments in which the antigen can bind.

Additionally, we observe that hot spot formation on both sides of the interface is particularly enriched by Trp and Tyr residues, underscoring the key role of aromatic side chains with some amphiphilic character in antibody design. Furthermore, we find that when strong interface hot spots are detected, they tend to persist in the apo conformation, suggesting that there is an inherent structural stability that surrounds core interface hot spots. These findings demonstrate the utility of computational solvent mapping for analyzing protein-protein interfaces, and highlights that at least in most cases antibodies drive antibody-antigen interactions.

**Statement of Significance:** Antibodies represent an important and expanding class of therapeutics for a range of diseases including infectious diseases, cancers, and autoimmune disorders. Enhancing our fundamental understanding of what drives antibody-antigen interactions is critical to our enhancing our ability to modulate these interactions. This work presents a systematic, physics-based study of antibody-antigen interfaces, and identifies key drivers of binding.

## 1 Introduction

Antibody-based therapeutics have become one of the fastest-growing classes of modern medicines, with applications ranging from oncology and autoimmune disorders to infectious diseases.^1^ These biologics exploit the exquisite specificity of antibodies for their antigens, allowing targeted modulation of disease-related pathways while minimizing off-target effects. The clinical success of monoclonal antibodies underscores their versatility as well as the desire to understand the molecular principles that govern antibody–antigen recognition.

The notion of binding “hot spots” has been widely accepted as a central concept in the study of biomolecular interactions. Broadly speaking, hot spots can be defined as regions on the receptor surface that disproportionately contribute to the binding free energy, and often act as the primary anchors of molecular recognition.^2^ Hot spots can be identified experimentally as regions on the receptor that can bind a variety of small organic probe molecules.^3–5^ Since probe molecules typically bind with weak affinity, biophysical techniques must be used to identify hot spots via this approach. In the structure activity relationship (SAR) by nuclear magnetic resonance (NMR) approach, the number of probe molecules that bind to a given site can be directly correlated to the druggability of the site.^2,3^ In the multiple solvent crystal structures (MSCS) method, the protein crystal is soaked in a variety of organic solvents, and solved multiple times in each solvent. Then, the structures are superimposed to identify hot spots as regions where a consensus of diverse solvent molecules bind.^4,5^

To expedite the speed and reduce the cost at which hot spots can be identified, a number of computational techniques for hot spot detection have been developed to mimic experimental approaches. These include the multiple copy simultaneous search (MCSS),^6–8^ mixed solvent molecular dynamics (MixMD),^9^ site identification by competitive ligand saturation (SILCS),^10^ and FTMap algorithm,^11,12^ which computationally sample interactions between the receptor and probe molecules to identify potential hot spots.

Despite the widespread use of therapeutic antibodies, there has to our knowledge been no comprehensive, systematic analysis of the hot spots at antibody–antigen interfaces using molecular probe-based hot spot detection methods. Because a relatively small number of such residues can dominate binding energetics, mapping them accurately is essential for rational antibody engineering, epitope selection for the design of immunodiagnostic assays or vaccines, and potentially predicting resistance mutations. Notably, the FTMap algorithm implements a fast Fourier transform (FFT) based mapping approach to identify hot spots as consensus sites (CSs), energy minima in which a variety of small organic probe molecules are predicted to bind.^11,12^ The consensus approach implemented in FTMap reduces false positives, and the implementation of an FFT-based algorithm significantly reduces the computational cost of the calculation. In the past, FTMap was successfully applied towards identifying protein–protein interfaces that were druggable by small molecules.^13^ In this work, we aim to apply FTMap to analyze hot spot formation at antibody–antigen interfaces, which are not conventionally druggable via small molecule inhibition. Instead, the larger aim of this work is to provide fundamental knowledge that will be useful for potentially designing improved antibody-antigen interfaces.

In this study, we assemble a curated dataset of high-resolution experimentally determined antibody–antigen structures taken from the publicly available ABAG-Docking benchmark set.^14,15^ We characterize hot spots that appear on the epitope and paratope via the FTMap algorithm, and identify residue types that appear to be enriched at antibody–antigen interfaces. Furthermore, we evaluate where complimentary hot spots appear in antibody–antigen complexes, and assess if interface hot spots can be identified by mapping the *apo* conformations of the subunits in antibody–antigen complexes. The following sections of this paper describe our data set in detail and present the analytical methods and key findings that reveal conserved principles of hot-spot formation at antibody-antigen interfaces, offering insights that could guide next-generation antibody design

## 2 Methods

### Dataset Construction

In this study, we analyzed the subset of 50 structures from the ABAG-Docking Benchmark with a resolution ≤ 3.0 Å.^14^ The extracted dataset is comprised of 50 antibody–antigen complexes, which we refer to in this work as the “bound set”. Additionally, we compile a corresponding “unbound set” within the resolution cutoff of 25 *apo* antibody structures for which an *apo* structure of the paired antigen is also available. A full list of the PDB IDs of the structures in the bound and unbound sets is available in the **Supplemental Information (Table S1–S2)**.

To establish ground-truth definitions for the epitope and paratope residues in each complex, we applied the following data analysis pipeline to each antibody–antigen complex in the bound set. For a given subunit in the complex, all residues within 10 Å of the binding partner are selected in PyMOL. For each residue in the selection, we calculate the solvent accessible surface area (SASA) in both the presence and absence of the binding partner using the get_sasa_relative() function in PyMOL. If the change in relative SASA for a residue is < 10%, it is removed from consideration. All remaining residues in the selection are then labeled as the paratope (if on the antibody) or epitope (if on the antigen) in a given structure. For structures in the unbound set, all equivalent residues are labeled as being part of the epitope or paratope. With this approach, we identify epitope and paratope selections that are consistent with previous analyses (see **Supplemental Information**).^16,17^

### Hot Spot Mapping with FTMap

To identify the locations of hot spots at antibody–antigen interfaces, we apply the FTMap hot spot mapping algorithm. FTMap has been applied extensively towards druggability analyses, and has been described in detail in previous publications.^11–13,18^ In short, FTMap applies a Fast Fourier Transform (FFT) docking approach to rapidly evaluate millions of potential interactions between a given receptor structure and 16 small molecular probes. For each probe type, the top 2000 best scoring binding poses are retained and energy minimized. Then, the minimized poses are clustered on the basis of their positions using a simple greedy algorithm. Finally, hot spots are identified as consensus sites–regions on the receptor surface where several probe clusters overlap. Consensus sites are ranked on the basis of their size, where consensus sites containing more probe clusters represent stronger hot spots. Previously, FTMap has been used to evaluate the druggability of hot spots on the protein surface by organic compounds. By our previously established definition, consensus sites with ≥ 16 probe clusters denote a hot spot that is likely to be druggable by a small molecule inhibitor, and consensus sites with a size 13–16 probe clusters are considered to be borderline druggable by small molecule inhibitors or peptides.^18^

For each antibody–antigen complex in the dataset, we apply FTMap to each subunit in isolation. For antibody subunits, we only map the variable (Fv) region with FTMap. This was done to focus the mapping calculation on antibody domains that were relevant for antigen binding. To automatically select Fv regions, the BioPython implementation of the DSSP algorithm was used to identify secondary structures in each antibody.^19,20^ Joints between the variable and constant region of the antibody are identified as loops without secondary structures. These loops are visually inspected, and then cut along their central residue to produce a structure containing only the Fv region.

### Characterizing Hot Spots and Interfaces

To identify FTMap hot spots that appear at the antibody–antigen interface, a set of interface inclusion criteria was determined. For the bound set, an antibody hot spot was considered to be at the paratope if it was within 3.5 Å of the paratope and within 6.0 Å of the epitope. Similarly, an antigen hot spot was considered to be at the epitope if it was within 3.5 Å of the epitope and within 6.0 Å of the paratope. For the unbound set, the corresponding bound structure was superimposed with the mapped subunit, and then the inclusion criteria were applied to select interface hot spots. In this work, we refer to an interface residue as a “hot spot residue” if it forms non-bonded interactions within 3.5 Å of any probes in an FTMap consensus site that appears at the interface.

Additionally, we characterize complimentary hot spots that appear at antibody–antigen interfaces. In this study, an FTMap consensus site is deemed to be a “complimentary” hot spot if it overlaps within 2.0 Å of an interface residue on the binding partner. By this definition, if an epitope residue binds within 2.0 Å of an interface hot spot that was predicted by mapping the antibody, the hot spot is considered to be complimentary. This analysis mirrors previous applications of the FTMap algorithm, in which chemical groups that bind within the volumes of predicted hot spots are presumed to be of greater importance for binding.

### Calculating the relative frequency of interface hot spot residues

To quantify if certain residue types were enriched as hot spot residues at the interface, we apply the following protocol. For a given residue type *i*, we calculate a normalized population value using the following equation:

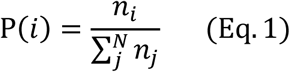

where *n_i_* is the number of times residue type *i* is observed as an interface residue in the benchmark set, and *n_j_* is the number of times residue type *j* is observed as a ground-truth interface residue in the benchmark set, and *N*=20, representing each of the 20 canonical amino acid types.

Normalized population values were calculated for ground-truth paratope and epitope residues, as well for the subset of hot spot residues identified at each interface. Then, the relative frequency of each residue type as a hot spot residue was calculated with the following approach:

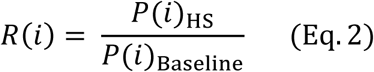

Where P(i)_HS_ is the normalized population for interface hot spot residues of type *i*, and P(i)_Baseline_ is the normalized baseline population for residues of type *i*. In this work, we use two different baseline populations to calculate relative frequency. This first metric, R(i)_any_, treats P(i)_Baseline_ as the population of any solvent-exposed residue *i*, across the entire antibody or antigen surface. We define a residue to be solvent-exposed if it is calculated to have a relative SASA value ≥ 25%. Relative SASA values are calculated using the crystal structure of the holo antibody–antigen complex. Separate *P(i)*_Baseline_ values are calculated for the antibody and the antigen. The second metric, R(i)_Interface_, is calculated using the *P(i)*_Baseline_ values that represent ground-truth interface residue populations. Based on these metrics, an R(*i*) > 1.0 indicates that residue type *i* will typically interact with an FTMap hot spot when it appears in the baseline population, and an R(*i*) < 1.0 suggests that residue type *i* is generally not observed to interact with FTMap hot spots.

### Assessing Hot Spot Conservation in Apo Structures

To assess if interface hot spots identified from mapping holo structure remain conserved in apo conformations, we apply FTMap to the unbound set of 25 structures described above. Apo structures were mapped and subsequently superimposed onto their holo counterparts with probe clusters using the align() function in PyMOL. Next, we select interface hot spots that were identified by mapping the holo conformation of the receptor. To measure the extent to which hot spot strengths change across states, we count the number of FTMap probe clusters predicted on the apo conformation that are placed within 1.0 Å of the initial holo consensus site. Then, we calculate the change in hot spot strength as the difference in size (number of probe clusters) between the original holo FTMap hot spot, and the number of overlapping FTMap probes predicted to bind on the apo structure. We opted to use this approach to compare changes in hot spot signal within the same overlapping volume on the receptor surface, which can more accurately reflect changes in hot spot strength that can result from conformational changes.

## 3 Results and Discussion

### Characterizing hot spots at antibody–antigen interfaces

In principle, hot spots appearing at the antibody–antigen binding interface disproportionately contribute to the binding free energy of the complex.^13,21,22^ In protein binding sites, hot spots typically appear on a small subset of the binding surface.^13^ Previously, we identified that druggable sites on protein–protein interfaces can be characterized as clusters of binding hot spots that possess a general tendency to bind side chains of the partner protein.^13^ Therefore, characterizing the locations, strengths, and frequencies at which hot spots appear in antibody–antigen interfaces can provide insights into the drivers of antibody–antigen recognition. In the first stage of our analysis, we aimed to characterize where hot spots appear at antibody–antigen interfaces by applying the FTMap algorithm to 50 holo antibody–antigen complexes deposited in the PDB (see **Methods**). All hot spot analyses discussed in this section were performed separately on the antibody and antigen subunits of a given complex (see **Methods**). In this work, we refer to the subset interface residues that form nonbonded interactions with FTMap probes as “hot spot residues” (see **Methods**, **Figure 1**).

**Figure 1.**
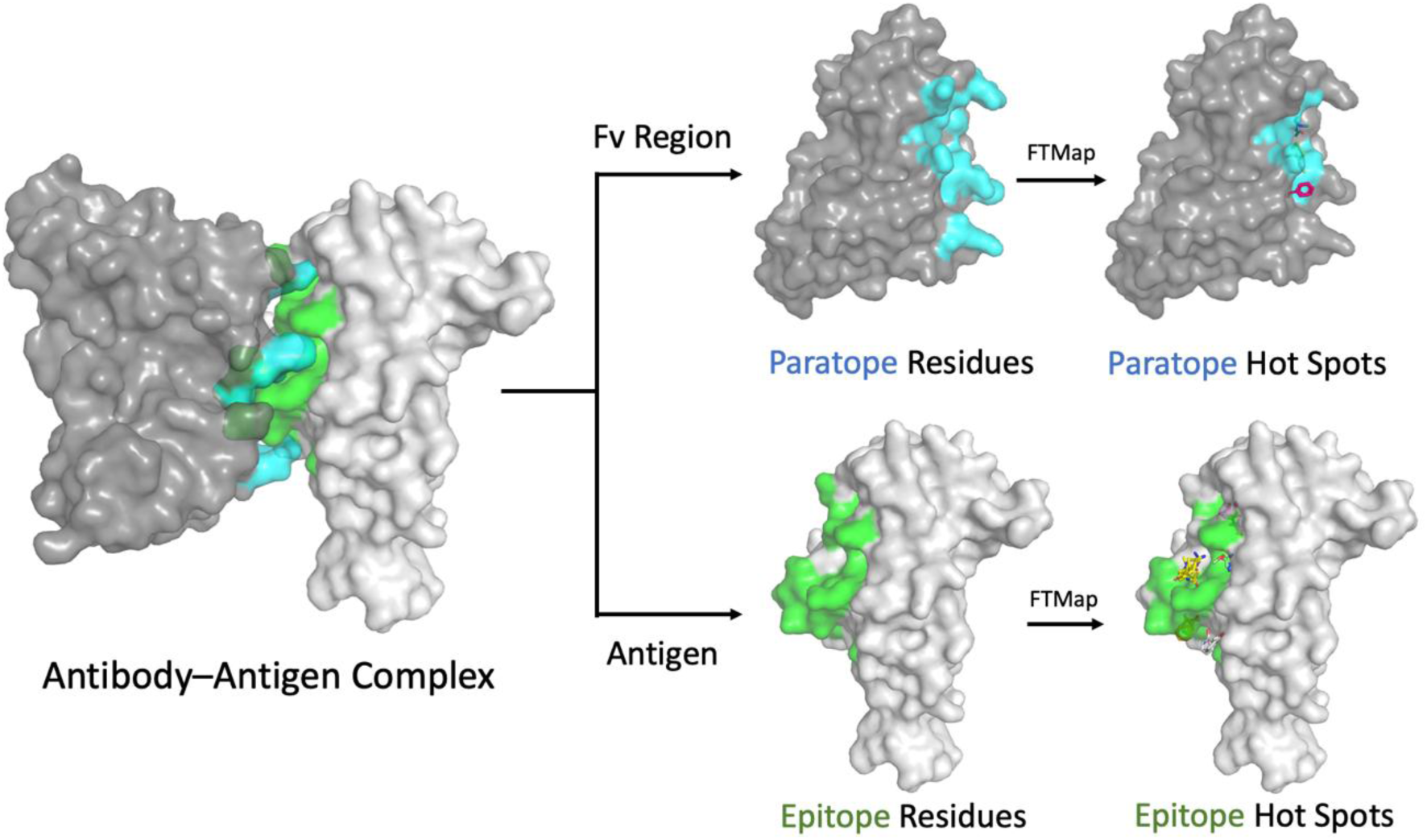
Hot spot mapping workflow. Paratope and epitope residues are identified from the holo Antibody–Antigen complex. Then, hot spots are identified on the Fv region and antigen chain using FTMap. Interface residues within 3.5Å of an FTMap consensus site are labeled as hot spot residues. The protein is shown in a grey surface representation, paratope residues are highlighted in blue, epitope residues are highlighted in green, and FTMap probes are shown in stick representation.

From our mapping calculations, we find that FTMap typically identifies more hot spot residues on the paratope side of the interface as opposed to the epitope side. Hot spots were identified at the paratope in 44/50 bound antibody structures and in 28/50 bound antigen structures. Excluding paratopes and epitopes with no predicted hot spots, the average number of consensus sites at the paratope and epitope were 2.3 and 1.8, respectively. (**Figure S2**). Additionally, we find that on average 34.7% of the residues in a given paratope are identified as hot spot residues, while an average of 14.1% of the residues in a given epitope are classified as hot spot residues (**Figure S3**).

Furthermore, the median paratope and epitope hot spot strengths were 8 and 8.5, respectively (**Figure S5**). Our findings support the notion that paratopes are hot spot rich environments that are primarily responsible for driving antibody–antigen binding.

In theory, if certain residue types are more likely to drive hot spot formation at the interface they are more likely to contribute disproportionately to the binding free energy of antibody–antigen complexes. Therefore, we aimed to measure the relative frequency of different amino acids at hot spots across all 50 structures in the bound set. To achieve this, we analyze the populations of all interface residues in the benchmark set, and compare them to the populations of hot spot residues that appear in the interface. We primarily use relative frequency, R(*i*), as a metric to identify residue types that are likely to be enriched as hot spots at the interface, and also to identify residues that are less likely to function as hot spots (see **Methods**).

Among all surface residues in antibodies in the bound set, we find that Ser, Gly, Thr, Lys, and Gln appear most often. When compared to the populations at interface residues, we observe that Gln, Lys, and Pro all experience large drops in population, suggesting that these amino acids are unlikely to appear as part of the paratope even though they appear on the antibody surface (**Figure 2a**, **Table S3**). Additionally, we find that Ser retains its place as the most probable solvent-exposed residue and most probable interface residue. Consistent with previous observations, Ser, Tyr, and Gly residues are frequently observed within the paratope (**Figure 2a**).^23–25^ Within the bound benchmark set, each of these residue types was found to have normalized interface populations of 0.20, 0.17, and 0.10, respectively (**Table S3**). Indicating that there is a preference for these residues to be included in the paratope. However, populations of these same amino acids among the subset of hot spot residues differs, suggesting that certain residue types may make larger contributions to the binding free energy despite appearing less often at the interface. Indeed, when calculating the relative frequency of paratope hot spot residues, we find that nine residues were observed to have an R(*i*)_interface_ > 1.0 (**Figure 2b**, see **Methods**). Among these residues the amino acids that have the largest interface relative frequencies are: Trp (R(i)_interface_ = 2.21), Arg (R(i)_interface_ = 1.66), Leu (R(i)_interface_ = 1.50), His, (R(i)_interface_ = 1.20), and Tyr (R(i)_interface_ = 1.19). When considering relative frequency with respect to all surface residues, we find particularly large R(i)_any_ values are obtained for aromatic residues: Trp (R(i)_any_ = 24.92), Tyr (R(i)_any_ = 7.57 ), His (R(i)_any_ = 4.12), and Phe (R(i)_any_ = 3.88) (**Figure 2b**, **Table S5**). Importantly, these residue types appear infrequently as solvent-exposed residues on the antibody, supporting the notion that the aromatic character of these amino acids is of particular importance in the paratope because they disproportionately contribute to hot spot formation.^26^

**Figure 2.**
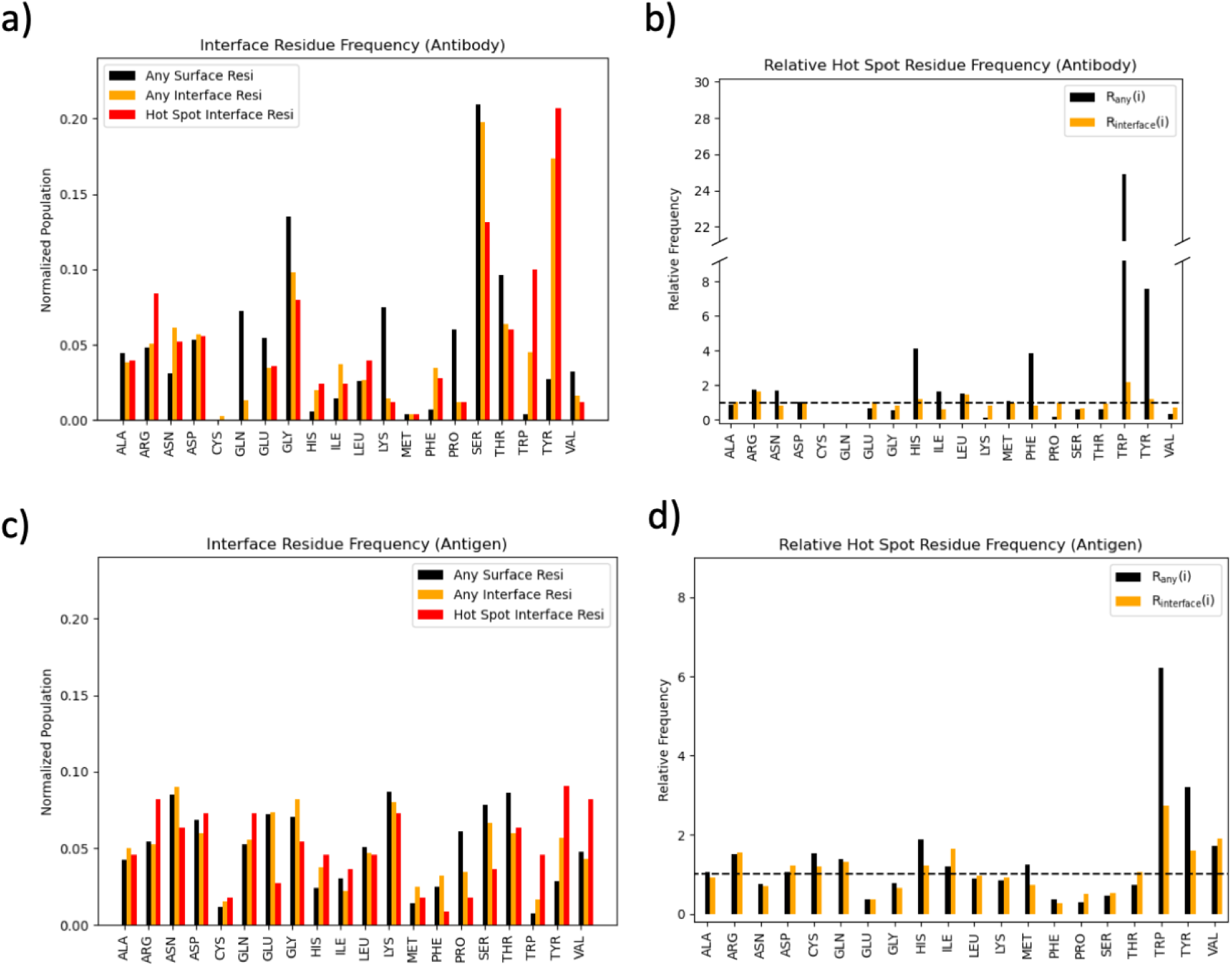
Residue population statistics across all structures in the bound set. a) Residue populations in the paratope, b) relative frequencies of hot spot residues in paratopes, c) residue populations in the epitope, and d) relative frequencies of hot spot residues in epitopes. representation.

Interestingly, we find that Ser and Gly have R(i)_interface_ and R(i)_any_ values < 1.0, suggesting that that although these residues appear often as hot spot residues in the bound benchmark set (**Figure 2a**), they do not disproportionately contribute to hot spot formation at the interface. We note that Ser and Gly are most frequently observed as surface residues and in CDR regions of antibodies in our benchmark (**Figure S2-S3**). This suggests that although these residues are an important component of the CDR loops, they are typically not directly contributing to the most energetically favorable interactions at the antibody–antigen interfaces in the subset of complexes analyzed in this study. It may be that Gly and to some extent Ser are present to allow these regions to be highly flexible^27^ One caveat of this observation is that the FTMap algorithm only identifies hot spots formed through direct protein–ligand interactions, and therefore does not identify residues that may impact the binding free energy through solvent-mediated interactions or entropic contributions. As a result, CDR residues with low R(i) values but high populations may still contribute to epitope recognition, albeit through mechanisms that FTMap is not designed to detect.

On the epitope side, the identification of residue types that are more likely to function as hot spot residues may provide insights that can assist with epitope prediction. In our analysis of all epitopes in the bound benchmark set, we find that the distribution of normalized populations for all residue types is relatively flat when compared to the results obtained from our paratope analysis (**Figure 2c**, **Table S4**). In general, we find that R(i)_any_ values on antigens do not reach the same magnitude as those observed for the most enriched residues on antibodies, with the largest relative frequencies being observed for: Trp (R(i)_any_ = 6.22), and Tyr (R(i)_any_ = 3.19). When comparing the relative frequencies of hot spot epitope residues with general epitope residues, we find the largest interface relative frequencies for Trp (R(i)_interface_ = 2.73), Val (R(i)_interface_ = 1.90), Ile (R(i)_interface_ = 1.64), Tyr (R(i)_interface_ = 1.60), Arg (R(i)_interface_ = 1.55), Gln (R(i)_interface_ = 1.31), Asp (R(i)_interface_ = 1.22), His (R(i)_interface_ = 1.21), and Cys (R(i)_interface_ = 1.19) (**Figure 2d**, **Table S5**). This suggests that these residues are more likely to contribute to the binding interface as hot spots, albeit not as disproportionately as the residues observed on antibody structures. Conversely, we see that Phe (R(i)_interface_ = 0.28), Glu (R(i)_interface_ = 0.37), Pro (R(i)_interface_ = 0.52), and Ser (R(i)_interface_ = 0.55), suggesting that these amino acids rarely contribute to hot spot formation at the antibody-antigen interface.

Based on our relative frequency analysis of antibody–antigen interfaces, we find that Trp and Tyr emerge as prominent hot spot residues on both sides of the interface. On the epitope side of the interface, we find that a broader range of residue types are observed to have R(i)_interface_ ≥ 1.10. These residues contain a mix of polar, hydrophobic, and charged side chains, indicating that epitopes tend to be amphiphilic regions on the receptor surface. For paratopes, we find high R(i) values for a smaller subset of residues with hydrophobic character or positively ionizable side chains. This is consistent with our observations of enriched hot spot residues within the paratope, and suggests that binding is primarily being driven by the antibody side of the interface, while epitope residues may contribute more to specificity.^28^ Additionally, we have previously noted that hot spots can be characterized as hydrophobic regions on the receptor surface that are decorated by patches of polarity.^29,30^ The disproportionate contributions of Trp and Tyr on both sides of the interface support the importance of this mosaic pattern, as these residues contains side chains with significant hydrophobic character in addition to including polar atoms. Thus, the arrangement of these residues in hot spots likely contribute some to specificity as well as binding.^31^ Furthermore, although a larger portion of the paratope is comprised of hot spot residues than the epitope, the appearance of relatively few hot spot residues at antibody–antigen interfaces reinforces the notion that antibody affinity is dictated by a handful of residue side chains.^32,33^

### Analyzing complimentary hot spots at antibody–antigen interfaces

In the context of small molecule binding, moieties that bind within hot spots anchor ligand binding, and therefore remain conserved as a ligand is optimized.^34^ Presumably, hot spots that appear at protein–protein interfaces perform a similar function, and help anchor the binding partner to the interface. In the context of antibody–antigen interfaces, we aimed to further characterize how complex formation is facilitated by performing an assessment of “complimentary” hot spots in antibody–antigen interfaces. In contrast to the analysis in the prior section which focused on identifying residues that drive hot spot formation, our complimentary hot spot analysis aims to characterize the locations and strengths of FTMap consensus sites in which residues from the partner protein bind.

Our protocol for performing complimentary hot spot analysis is described in the **Methods** section. In short, FTMap results were obtained for each subunit of each complex in the bound benchmark set are superimposed with their binding partner. Then, the subset of FTMap consensus sites that overlap within 2.0 Å of a residue on the binding partner are selected as complimentary hot spots. By this definition, complimentary hot spots that appear on the epitope are defined as consensus sites within which paratope residues bind, and visa-versa. A graphical overview of the complementary hot spot analysis protocol is displayed in **Figure S6**.

When considering hot spots of any strength, we find that at least one complimentary hot spot is identified by FTMap in 80% of the structures in the bound benchmark set. As displayed in **Figure 3a**, a further breakdown of these structures reveals that complimentary hot spots appear on the epitope in only 10% of cases, on the paratope only in 48% of cases, and on both sides of the interface in 22% of cases. Notably, complimentary hot spots appear on the epitope in 32% of cases in the bound benchmarks set, indicating that it is rare for epitopes to be identified via conventional hot spot-based analyses. Conversely, complimentary hot spots appear on the paratope in 70% of the mapped complexes. Coupled with our previous observation that a larger fraction of the paratope is covered in hot spots than epitope counterparts (**Figure S4**), our analysis highlights that paratopes act as hot spot-rich environments within which the antigen can easily bind. In a future study, we aim to further investigate the relationship between antibody maturation and hot spot formation to further understand the role of binding hot spots in high-affinity antibody binding.

**Figure 3.**
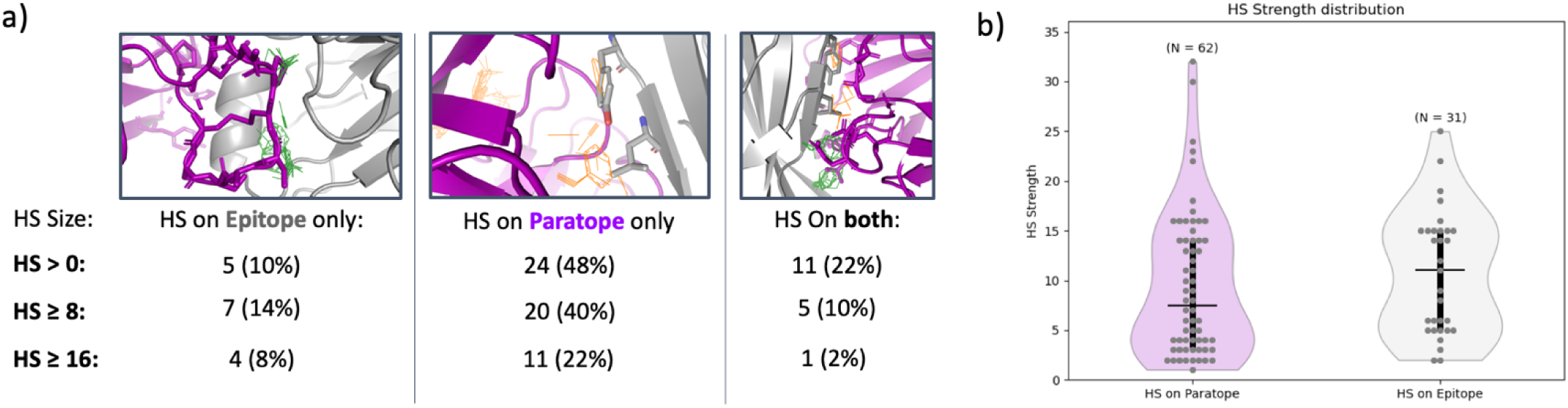
Overview of complimentary hot spot analysis. a) A table summarizing the number and percentage of cases in the bound benchmark set where complimentary hot spots are observed on the epitope only, the paratope only, or on both sides of the interface. Results are displayed using minimum hot spot strength cutoffs of > 0, ≥ 8, and ≥ 16. The antigen is displayed in grey, the antibody is displayed in purple, antigen hot spots are shown as green sticks, and antibody hot spots are shown as orange sticks. b) Violin plots displaying the distribution of complimentary hot spot strengths observed on the paratope and epitope. The total number of observed complimentary hot spots is listed above each respective violin plot. The median value is denoted by a horizontal black line, interquartile ranges are shown as vertical black bars, and each data point is plotted as a grey dot.

Additionally, we examine the strengths of complimentary hot spots that were identified in our analysis. When applying FTMap to assess the ligandability of small molecule binding sites, a hot spot strength of 16 or greater typically denotes a druggable site.

However, if we only consider complimentary hot spots with a strength of 16 or larger, we find that these sites are only observed in 32% of the complexes in the test set (**Figure 3a**). Furthermore, only 10% of the cases in the benchmark set were found to have druggable complimentary hot spots on the epitope, only one of which was identified to have druggable hot spots on both sides of the interface. When viewing the distributions of complimentary hot spot strengths, we find that although complimentary hot spots appear more often on the paratope side of the interface, they are typically weaker than those that form on the epitope (**Figure 3b**). On paratopes where hot spots are predicted, a median of 2 consensus sites appear, with a median strength of 8 probe clusters. Whereas on epitopes where hot spots are identified, a median of 1 consensus site is observed, with a median strength of 11 probe clusters. This discrepancy in median hot spot strengths can be attributed to the fact that weak consensus sites detected on the paratope are often accompanied by at least one stronger hot spot. (**Figure S7**). In contrast, single complimentary hot spots are more frequently observed on the epitope side of the interface.

These results indicate that it is rare to observe canonically druggable hot spots at antibody–antigen interfaces, and supports the notion that these interfaces cannot typically be targeted by small molecule inhibitors. For peptides binding to proteins the hot spot strength is typically ≥ 13,^18^ we show herein that for proteins binding to proteins at least for antibody antigen interactions the strength of the top-ranking hot spot at the interface is typically lower than that. As described above, this lower hot spot strength may be compensated for by the additional lower ranking hot spots. Furthermore, the paucity of strong hot spots on the epitope emphasizes the fact that antibodies bind to antigen interfaces that are traditionally not druggable by more traditional small molecule binders.

### Comparison of Hot Spot Mapping Between Bound and Unbound Structures

In principle if hot spot mapping can be used to identify epitope residues on apo conformations of the antigens, the approach may be useful in prospective applications, such as epitope prediction and the rapid design of immunodiagnostic assays.

Furthermore, this analysis may highlight limitations of FTMap in this context, and lay the groundwork for future improvements that can be made to the methodology within the context of epitope detection and paratope characterization.

In order to investigate if hot spots are conserved in the apo conformations of the antibody and antigen, we compare FTMap results between the bound and unbound set of antibody–antigen structures. In order to compare results, we map a subset of 25 cases in the ABAG docking set which contained apo and holo structures that satisfied our 3.0 Å resolution cutoff (see **Methods**).

First, we evaluated if the strength of interface hot spots varied between the apo and holo conformations of the subunits in each complex. In the ABAG-Docking benchmark set, structures are categorized into three categories on the basis of their conformational flexibility.^14^ Targets are classified as “rigid”, “medium”, or “difficult” based on a number of statistical criteria described by Zhao et al.^14^ Rigid targets undergo minimal conformational changes upon binding, whereas difficult targets are observed to have larger conformational reorientations at the binding interface. In the unbound set used in this work, there are 11 rigid complexes, 8 medium complexes, and 6 difficult complexes. For antibodies, we find that as the flexibility of the paratope increases, the median hot spot strengths obtained by FTMap generally decrease. Comparing bound and unbound paratope conformations, we observe median strengths of 11 and 13, 9.5 and 12, and 7 and 7 for the rigid, medium, and difficult cases, respectively (**Figure 4**). On the epitope side, we see a similar trend, with median hot spot strengths of 11.5 and 12, 6 and 9, and 7 and 10 on the bound and unbound structures for rigid, medium, and difficult cases, respectively.

**Figure 4.**
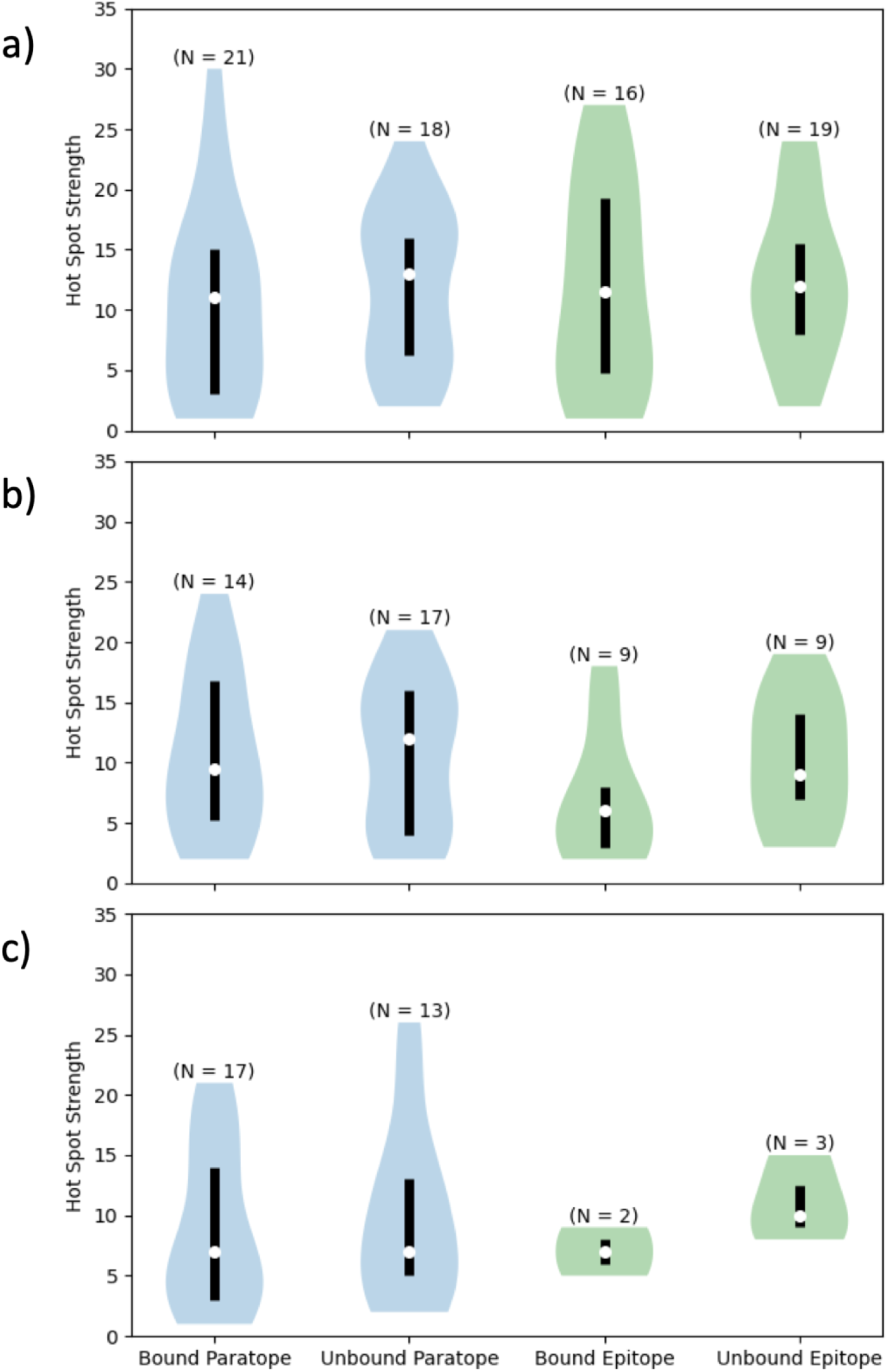
Distribution of FTMap hot spot strengths observed in on the bound paratope, unbound paratope, bound epitope, and unbound epitope. Results are split by structure flexibility, using the ABAG-Docking Benchmark designations of a) rigid, b) medium, and c) difficult. Number of hot spots in each distribution are displayed above each violin plot.

Across all targets, we find that on both the epitope and the paratope, interface hot spot strengths increase in the unbound structures when compared to their holo counterparts (**Figure S8**). Notably, we see larger gains in hot spot strength on the epitope side of the interface, where the median epitope hot spot strength increased greatly from 6 in the bound structures to 12 in the unbound structures. In principle, these gaps in median hot spot strength are expected when using a molecular probe-based method such as FTMap. This is primarily because apo conformations of the antibody typically have larger pockets than their holo counterparts. As a result, a larger number of probe clusters can form at the interface, and a stronger signal can be obtained by the method. In the more flexible “difficult” cases, we find that hot spot strengths are typically lower, largely due to the fact that binding pocket are less likely to be pre-formed in the apo conformation. In future applications, it may be desirable to augment FTMap-based hot spot analysis with conformational sampling techniques and to use FTMove.^35,36^

Next, we analyzed whether the locations of interface hot spots in the unbound structure are conserved in the holo structure (see **Methods**). Among the cases where interface hot spots are detected, we find that they typically remain conserved across both apo and holo conformations (**Table 2**). Hot spots at the interface may be less stable because there are more conformational changes that happen at the interface than at other areas of the structures. Notably, stronger “borderline druggable” hot spots with scores (number of probe clusters) ≥ 13 are more likely to be conserved than weaker hot spots. This is consistent with our previous understanding of small molecule binding, in which a strong “primary” hot spot is typically conserved in the apo conformation of the receptor. Strong hot spots were more likely to be conserved, both across the whole structure and at the paratope and epitope.

**Table 2.** Percent of apo interface hot spots whose locations are conserved in the holo structure.

| | All Hot Spots | Hot Spots (Size $\geq 13$) |
| --- | --- | --- |
| Paratope | 63.6% | 77.8% |
| Epitope | 69.7% | 85.2% |

To further examine the extent to which hot spot strengths vary between bound and unbound structures, we calculate the change in hot spot size for interface hot spots observed on holo structures (see **Methods**). With this analysis, our aim was to assess how hot spot strengths change within the same overlapping volume on the receptor surface, which in principle reflects perturbations in hot spot strengths that result from conformational changes. Based on this analysis, we find that hot spots are typically weaker in holo paratope conformations for rigid and medium flexibility cases (**Figure S9**). In results obtained for the rigid case 6QFC, we observe that the paratope only undergoes minor side chain reorientations upon antigen binding. As a result, the pocket formed by the paratope is more open in the apo conformation, and can accommodate additional FTMap probes (**Figure 5a**). This is reflected in the highlighted hot spot sizes, in which the apo hot spot has a stronger signal of 27 probe clusters, a slight increase over the 23 probe clusters observed in the holo hot spot (**Figure 5a**). Thus, if the interface undergoes limited conformational changes, hot spots tend to remain detectable on apo structures.

**Figure 5.**
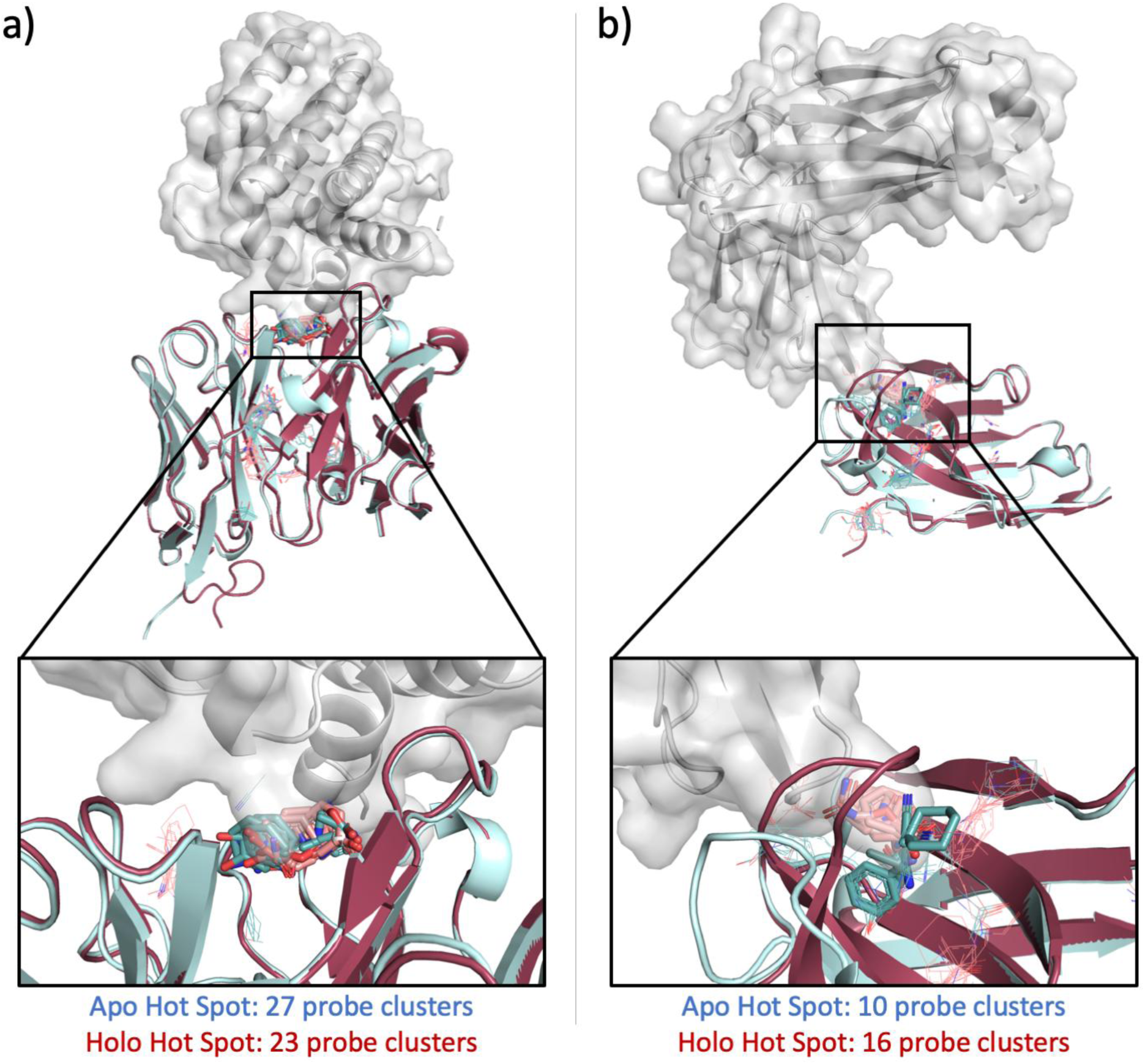
Conserved interface hot spots observed in apo conformations. Examples are shown for a) 6QFC, a rigid example, and b) 8F8X, a difficult example. The antigen is shown in grey, the apo antibody is shown in light blue, and the holo antibody is shown in magenta. FTMap probe clusters predicted on the apo antibody conformation are shown in teal line representation, and probe clusters predicted on the holo structure are shown in salmon. Conserved interface probe clusters are highlighted in stick representation.

In contrast, for “difficult” cases with high flexibility, we find that holo paratope hot spots are generally stronger than their apo counterparts, reflecting the fact that larger conformational reorientations can have a significant impact on hot spot formation at the interface. In **Figure 5b**, we highlight the difficult case 8F8X, in which a CDR loop is observed to undergo a large conformational change upon antigen binding. In this example, the holo conformation of the paratope is observed to clamp down around the antigen, creating a pocket in which a complimentary FTMap hot spot is observed to form. In the apo structure, the loop adopts an open conformation, creating a more broad binding site on the paratope. As a result, FTMap probes do not occupy the same volumes within the binding site, and a weaker consensus site is observed for the apo structure at the interface. Although there is a limited set of highly flexible structures in our analysis, this result emphasizes the fact that the conformational landscape of the paratope can impact the locations of hot spots.

## Conclusion

In this work, we apply the FTMap algorithm to a subset of 50 antibody–antigen complexes deposited in the ABAG-Docking benchmark set. From our analysis of holo structures, we find that hot spot formation primarily occurs on the paratope side of the interface.Trp, Tyr, Phe, and His are enriched as hot spot residues with respect to their baseline populations as interface residues. These findings indicate that these residue types may be particularly important for driving hot spot formation, and contribute towards creating a favorable binding environment within which epitope residues can bind. In addition, the paratope side often contains a larger number of weaker hot spots with multiple hot spots contributing to binding. Significantly, Trp and Tyr, both of which are aromatic residues containing polar atoms as well, are enriched as hot spot residues on both sides of the interface. However, we find that it is rare for hot spots to appear on the epitope side of the interface alone. This underscores the notion that epitopes are not typically druggable by small molecule therapeutics, and highlights why larger antibody-based therapeutics are needed to target these types of interfaces.

Through our analysis of apo structures, we find that when hot spots which are detected at the interface in apo structures are typically conserved in *holo* conformations. This is especially true when FTMap identifies stronger consensus sites. Additionally, we find that strong hot spot detection remains a challenge on highly flexible targets. To further improve the applicability of FTMap within the context of hot spot detection on apo antibody and antigen structures, we intend to explore how modifications to the scoring function, the expansion of the probe set, and the integration of conformational sampling can improve the performance.

## Data Availability

All structures in the ABAG-Docking benchmark set were originally published by Zhao et al.,^14^ and are available for download on GitHub: https://github.com/Zhaonan99/Antibody-antigen-complex-structure-benchmark-dataset/tree/master The FTMap algorithm is available as a free web server for academic and governmental users: https://ftmap.bu.edu/

## Author Contributions

O.K. and M.K. conducted the research and wrote the manuscripts. M.S. and H.C. conducted research. N.M. contributed code for data preparation. S.V. supervised the research. D.J.-M. planned and supervised the research and writing.

## Declaration of Interests

Sandor Vajda holds equity in Acpharis Inc, a company that developed the ATLAS program, a version of FTMap. Acpharis Inc offers commercial licenses for the use of ATLAS software; however, FTMap is freely available to academic and governmental users.

## Acknowledgements

The authors would like to thank Siri Allegra-Berger for helpful discussions and contributions to dataset selection. This work was supported in part by grant R35GM118078 from the National Institute of General Medical Sciences.

## Notes

### Competing Interest Statement

The authors have declared no competing interest.

