## Supplemental Tables S1-S8 and Figures S1-S9. for "Towards Understanding the Drivers of Antibody-Antigen Binding"

**[1] Department of Chemistry**

**[2] Department of Biomedical Engineering**

**Boston University**

**Boston, MA, USA**

**[#]These authors contributed equally**

**Keywords: Hot spot mapping, fragment positioning, antibody-antigen interfaces**

**Table S1.** PDB IDs for the subset of ABAG-Docking structures in the bound dataset.

| Structure Index | ABAG-Docking PDB ID |
|---|---|
| 1 | 6HER |
| 2 | 6HHD_1 |
| 3 | 6HHD_2 |
| 4 | 6JB8 |
| 5 | 6LGW |
| 6 | 6OTC |
| 7 | 6P4B |
| 8 | 6P50 |
| 9 | 6PXH |
| 10 | 6PZW |
| 11 | 6Q0O |
| 12 | 6QB6 |
| 13 | 6QFA |
| 14 | 6QFC |
| 15 | 6SV2 |
| 16 | 6U12 |
| 17 | 6U54 |
| 18 | 6XZF |
| 19 | 6YIO |
| 20 | 6YLA |
| 21 | 6Z3P |
| 22 | 6Z3Q |
| 23 | 6ZTR |
| 24 | 7C01 |
| 25 | 7CJ2 |
| 26 | 7CQC |
| 27 | 7DUO |
| 28 | 7KET |
| 29 | 7KF0 |
| 30 | 7KF1 |
| 31 | 7KFW |
| 32 | 7LVW |
| 33 | 7M3N |
| 34 | 7NP1 |
| 35 | 7NX3_1 |
| 36 | 7NX3_2 |

| | |
|---|---|
| 37 | 7R40 |
| 38 | 7SGM |
| 39 | 7SHY |
| 40 | 7SU0 |
| 41 | 7SU1 |
| 42 | 7T25 |
| 43 | 7TYV |
| 44 | 7VYR |
| 45 | 7WRV |
| 46 | 7Z2M |
| 47 | 8B7W |
| 48 | 8F8X |
| 49 | 8GV7 |
| 50 | 8GZ5 |

**Table S2.** PDB IDs for the subset of ABAG-Docking structures in the unbound dataset.

| Structure Index | ABAG-Docking PDB ID |
|---|---|
| 1 | 6JB8 |
| 2 | 6OTC |
| 3 | 6P4B |
| 4 | 6P50 |
| 5 | 6PXH |
| 6 | 6QB6 |
| 7 | 6QFC |
| 8 | 6U12 |
| 9 | 6U54 |
| 10 | 6XZF |
| 11 | 6ZTR |
| 12 | 7CJ2 |
| 13 | 7CQC |
| 14 | 7DUO |
| 15 | 7KF0 |
| 16 | 7KF1 |
| 17 | 7LVW |
| 18 | 7SGM |
| 19 | 7SU1 |
| 20 | 7T25 |
| 21 | 7Z2M |
| 22 | 8B7W |

| | |
|---|---|
| 23 | 8F8X |
| 24 | 8GV7 |
| 25 | 6UUH |

**Table S3**. Normalized populations for antibody residues

| Residue | Solvent-Exposed | Interface | Interface Hot Spot |
|---|---|---|---|
| ALA | 0.045 | 0.038 | 0.04 |
| ARG | 0.048 | 0.05 | 0.084 |
| ASN | 0.031 | 0.061 | 0.052 |
| ASP | 0.053 | 0.057 | 0.056 |
| CYS | 0 | 0.003 | 0 |
| GLN | 0.072 | 0.013 | 0 |
| GLU | 0.055 | 0.034 | 0.036 |
| GLY | 0.135 | 0.098 | 0.08 |
| HIS | 0.006 | 0.02 | 0.024 |
| ILE | 0.015 | 0.037 | 0.024 |
| LEU | 0.026 | 0.027 | 0.04 |
| LYS | 0.075 | 0.015 | 0.012 |
| MET | 0.004 | 0.004 | 0.004 |
| PHE | 0.007 | 0.034 | 0.028 |
| PRO | 0.06 | 0.012 | 0.012 |
| SER | 0.21 | 0.198 | 0.131 |
| THR | 0.096 | 0.064 | 0.06 |
| TRP | 0.004 | 0.045 | 0.1 |
| TYR | 0.027 | 0.174 | 0.207 |
| VAL | 0.032 | 0.016 | 0.012 |

**Table S4**. Normalized populations for antigen residues

| Residue | Solvent-Exposed | Interface | Interface Hot Spot |
|---|---|---|---|
| ALA | 0.043 | 0.05 | 0.045 |
| ARG | 0.054 | 0.053 | 0.082 |
| ASN | 0.085 | 0.09 | 0.064 |
| ASP | 0.069 | 0.06 | 0.073 |
| CYS | 0.012 | 0.015 | 0.018 |
| GLN | 0.053 | 0.055 | 0.073 |
| GLU | 0.072 | 0.074 | 0.027 |
| GLY | 0.071 | 0.082 | 0.055 |
| HIS | 0.024 | 0.037 | 0.045 |
| ILE | 0.03 | 0.022 | 0.036 |
| LEU | 0.05 | 0.047 | 0.045 |

| | | | |
|---|---|---|---|
| LYS | 0.087 | 0.08 | 0.073 |
| MET | 0.014 | 0.025 | 0.018 |
| PHE | 0.025 | 0.032 | 0.009 |
| PRO | 0.061 | 0.035 | 0.018 |
| SER | 0.079 | 0.067 | 0.036 |
| THR | 0.087 | 0.06 | 0.064 |
| TRP | 0.007 | 0.017 | 0.045 |
| TYR | 0.028 | 0.057 | 0.091 |
| VAL | 0.048 | 0.043 | 0.082 |

**Table S5.** Relative population values for antibody and antigen targets

| | **Antibody** | | **Antigen** | |
|---|---|---|---|---|
| **Residue** | $\mathbf{R_{Any}(i)}$ | $\mathbf{R_{Interface}(i)}$ | $\mathbf{R_{Any}(i)}$ | $\mathbf{R_{Interface}(i)}$ |
| ALA | 0.89 | 1.04 | 1.06 | 0.91 |
| ARG | 1.73 | 1.66 | 1.50 | 1.55 |
| ASN | 1.68 | 0.85 | 0.75 | 0.71 |
| ASP | 1.05 | 0.98 | 1.06 | 1.22 |
| CYS | 0.00 | 0.00 | 1.53 | 1.19 |
| GLN | 0.00 | 0.00 | 1.38 | 1.31 |
| GLU | 0.66 | 1.04 | 0.38 | 0.37 |
| GLY | 0.59 | 0.81 | 0.77 | 0.67 |
| HIS | 4.12 | 1.20 | 1.88 | 1.21 |
| ILE | 1.64 | 0.64 | 1.19 | 1.64 |
| LEU | 1.53 | 1.50 | 0.90 | 0.96 |
| LYS | 0.16 | 0.82 | 0.83 | 0.90 |
| MET | 1.11 | 1.00 | 1.26 | 0.73 |
| PHE | 3.88 | 0.81 | 0.37 | 0.28 |
| PRO | 0.20 | 1.00 | 0.30 | 0.52 |
| SER | 0.63 | 0.67 | 0.46 | 0.55 |
| THR | 0.62 | 0.94 | 0.73 | 1.07 |
| TRP | 24.92 | 2.21 | 6.22 | 2.73 |
| TYR | 7.57 | 1.19 | 3.19 | 1.60 |
| VAL | 0.37 | 0.75 | 1.72 | 1.90 |

Validation of ground-truth hot spot definitions:

We have determined the paratope and epitope from the bound set. To confirm that we have correctly identified the paratope and epitope, we observed that the average paratope consisted of 15.1 residues and the average epitope consisted of 14.4 residues. Previous literature describes paratopes and epitopes as typically between 12 to 16 residues long.[1] To further confirm that our results were characteristic of paratopes and epitopes, the distribution of residue types at the determined paratope and epitope was analyzed (**Figure S1**). Serine and tyrosine were found to be the most abundant residues at the paratope. Previous literature also observed a similar distribution of paratope residues.[2] Epitope residues were more uniformly distributed across residue types, with asparagine being the most abundant.

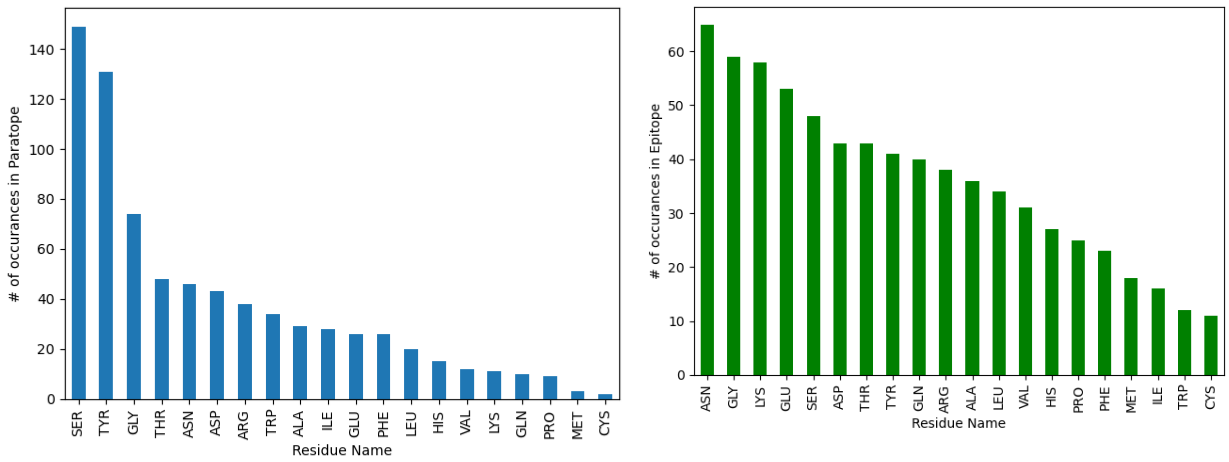


**Figure S1.** Left: Histogram of amino acid residues at paratope. Right: Histogram of amino acid residues at epitope.

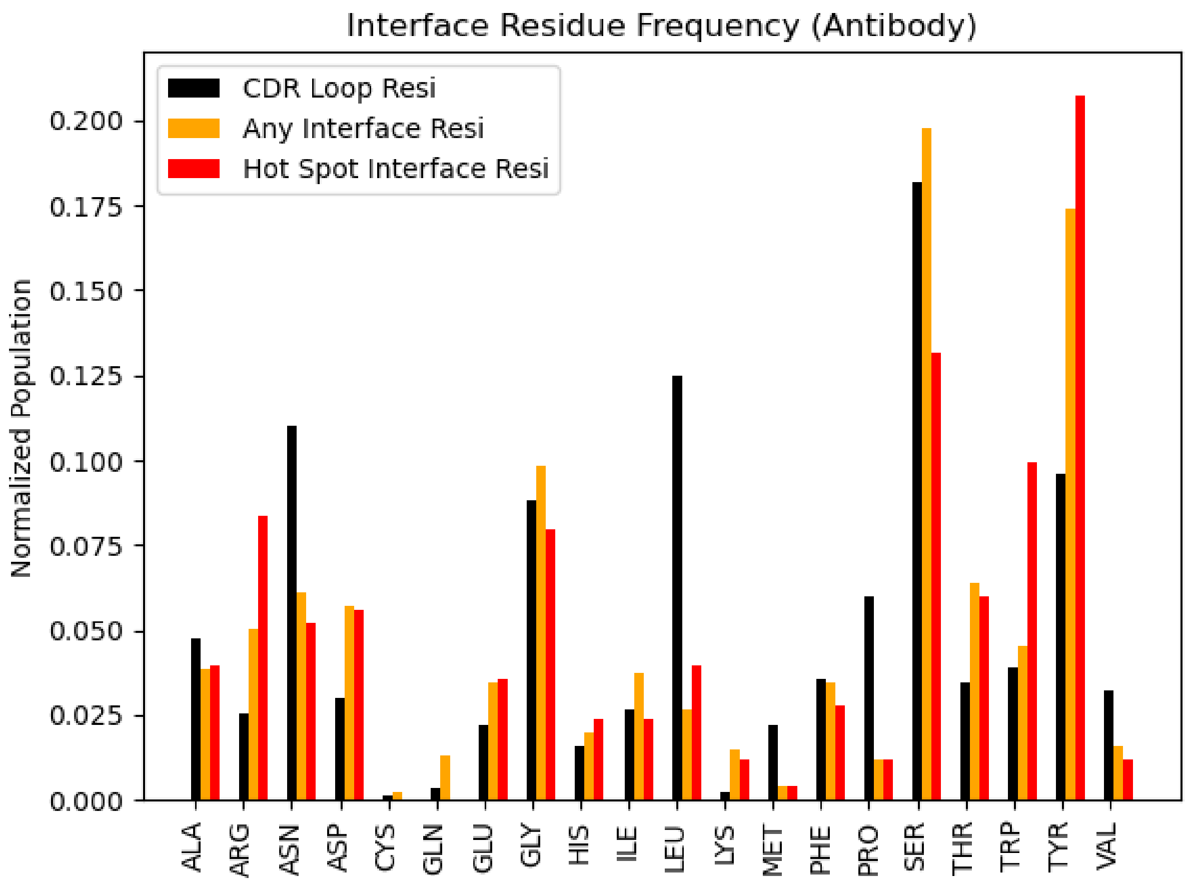


**Figure S2.** Populations of each residue type among CDR regions (black), interface residues (orange) and hot spot interface residues (red) across all antibodies in the bound benchmark set.

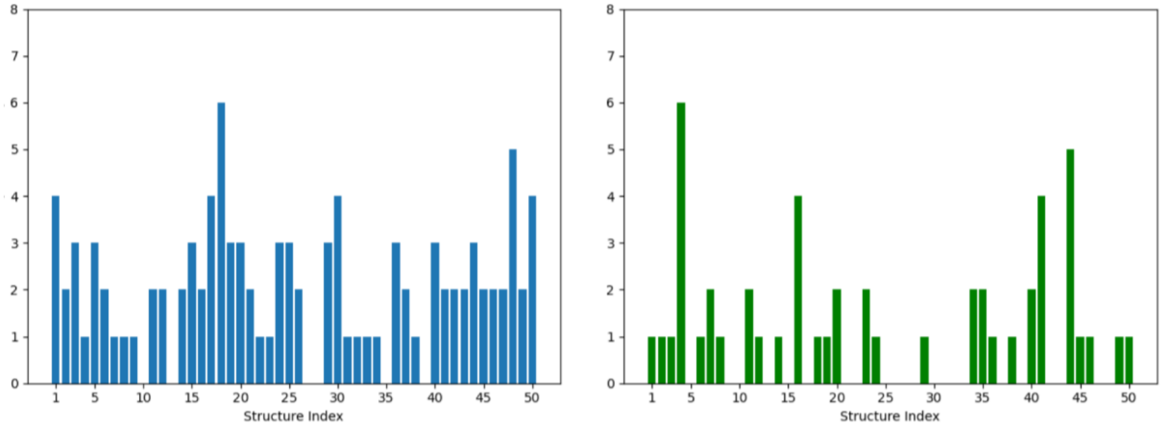


**Figure S3.** Number of FTMap consensus sites at each interface in the bound set. Left: Number of hot spots identified at paratope. Right: Number of hot spots identified at epitope.

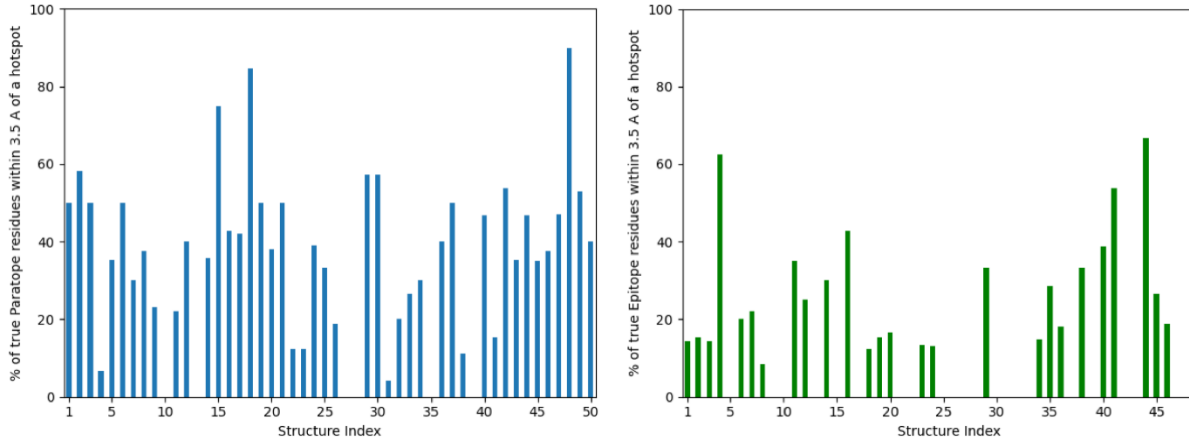


**Figure S4** – Left: Percentage of paratope residues that have a non-bonded interaction with a paratope hot spot. Right: Percentage of epitope residues that have a non-bonded interaction with an epitope hot spot.

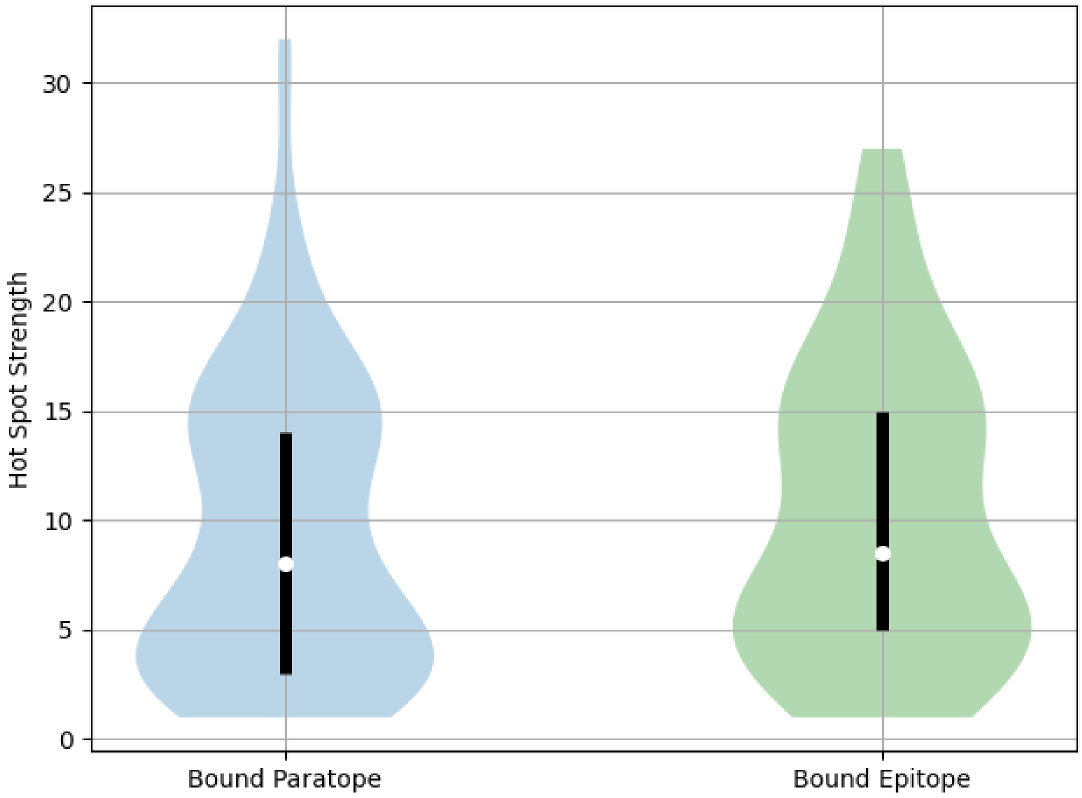


**Figure S5.** Distribution of FTMap hot spot strengths on paratope and epitope residues in the bound benchmark set. Median values are denoted by a white dot, and interquartile ranges are denoted by the black bars.

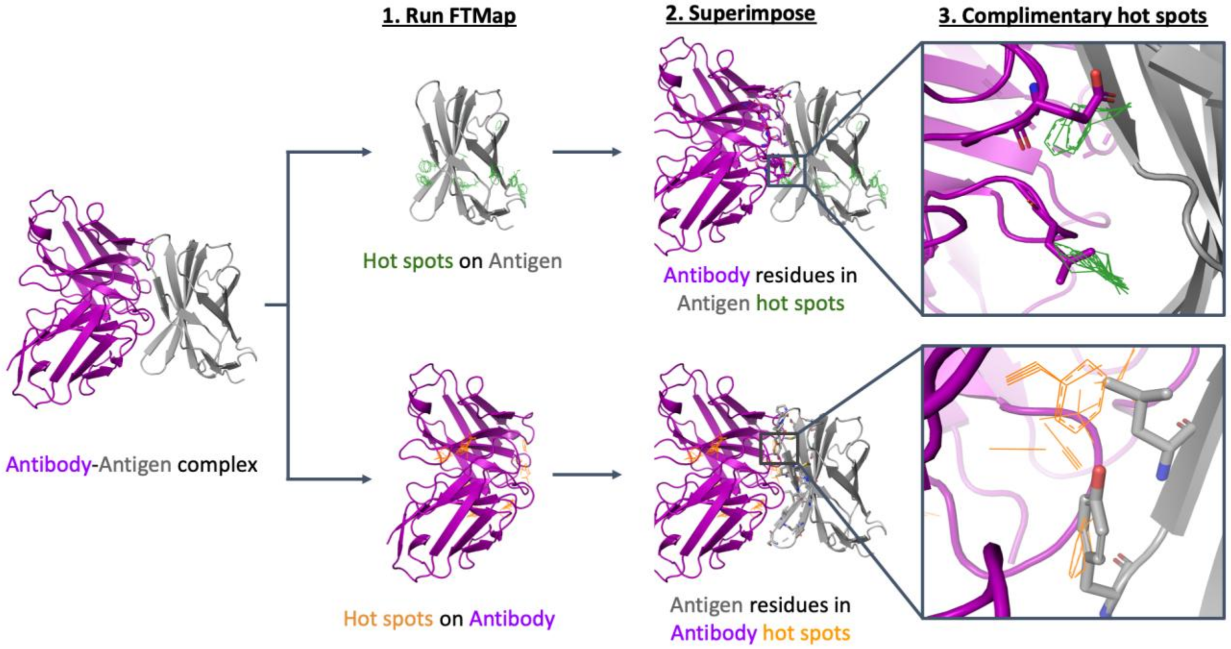


**Figure S6.** Graphical overview of the pipeline used to identify complimentary hot spots. First, FTMap is used to predict hot spots on the Antigen (grey cartoon) and antibody (purple cartoon) subunits separately. Next the binding partner is superimposed onto the mapped structure. Then, the subset of FTMap consensus sites (wire) that overlap within 2.0Å of a residue on the binding partner are selected as complimentary hot spots.

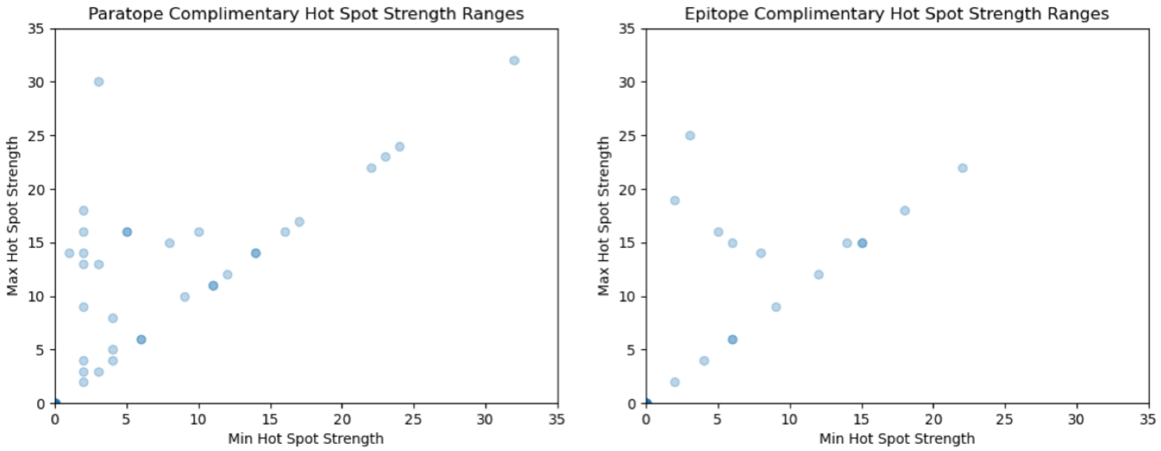


**Figure S7.** Maximum hot spot strength vs minimum hot spot strength for all cases in the bound benchmark set. The maximum and minimum hot spot strengths for a given structure in the bound benchmark set is displayed as a transparent blue dot. Points along the diagonal indicate that the maximum and minimum hot spot sizes are the same.

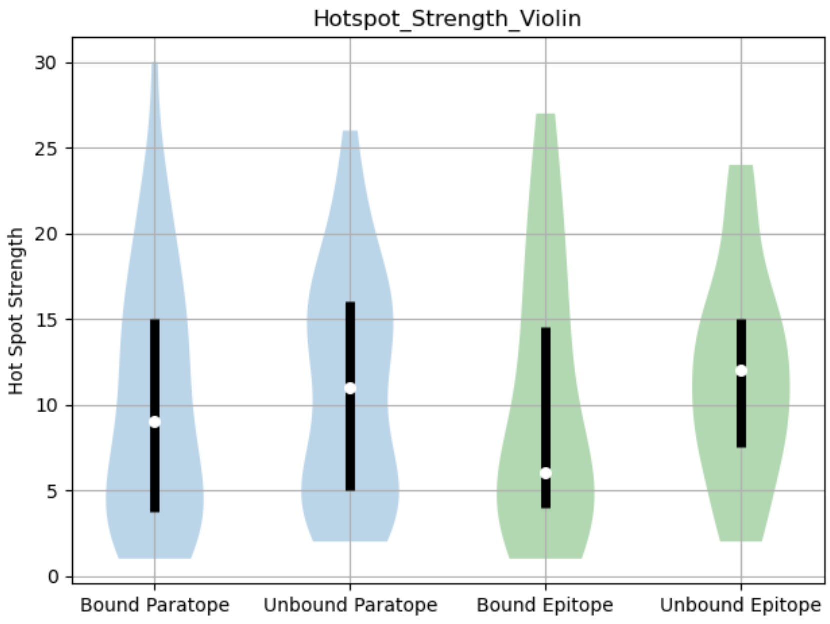


**Figure S8.** Violin plot of hot spot strengths observed on the bound paratope, unbound paratope, bound epitope, and unbound across all structure flexibilities. Median is displayed as a white dot. The interquartile range is shown as a black bar.

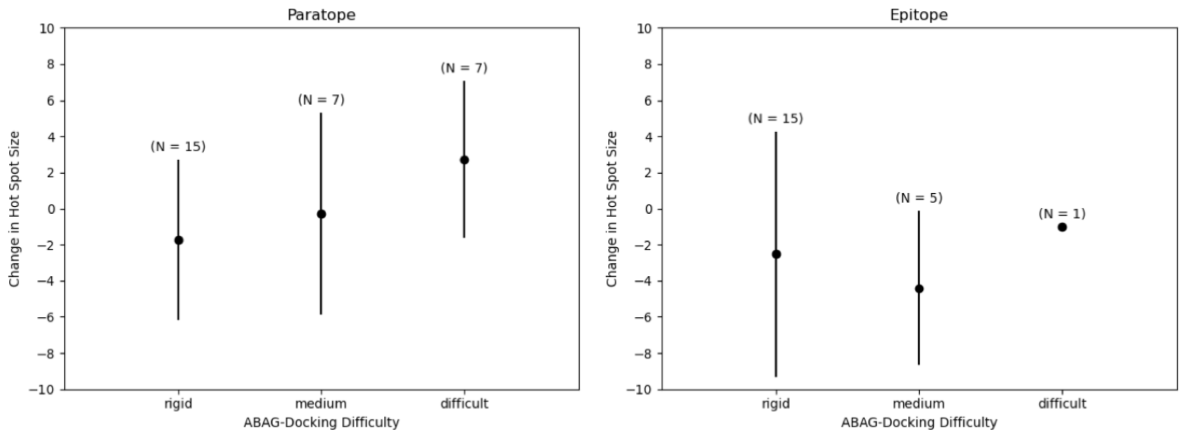


**Figure S9.** Average change in interface hot spot strengths between the bound and unbound conformations of the (Left) paratope and (Right) epitope. Standard deviation ranges are shown as black bars. Results are split by conformer flexibility, represented as ABAG-Docking Difficulty. The number of interface hot spots considered are annotated above each point. Negative values indicate that the holo conformation contains a weaker hot spot than the apo conformer, and positive values indicate that the holo conformation has a stronger hot spot than the apo variant.
